# Sclerotome-derived vascular smooth muscle progenitors contribute to the haematopoietic stem cell specification niche

**DOI:** 10.1101/2023.08.09.552695

**Authors:** Clair M. Kelley, Nicole O. Glenn, Dafne Gays, Massimo M. Santoro, Wilson K. Clements

## Abstract

Haematopoietic stem cells (HSCs) are the self-renewing progenitors that continuously populate the haemato-immune cell lineages throughout life, and constitute the therapeutic component of bone marrow transplants. A major biomedical goal has been to understand the native specification of HSCs during embryonic development as a means to inform *in vitro* directed differentiation of pluripotent stem cells. Across vertebrate phyla, HSCs derive from haemogenic endothelium in the ventral floor of the primitive dorsal aorta (DA), also known as the descending aorta in mammals. Competent HSC-fated cells in the endothelium likely receive instructive signaling from neighbouring cells that constitute a “specification niche.” We previously showed that experimental manipulations leading to defects in the most ventral compartment of the somite, the sclerotome, are correlated with HSC defects, raising the possibility that sclerotome patterning is required for HSC specification. Here we show that in zebrafish, specific sclerotome-derived cells contact the DA immediately prior to the emergence of HSCs. These cells subsequently give rise to vascular smooth muscle cells (VSMCs). When sclerotome patterning is disrupted, VSMCs are diminished, and HSC specification fails. We conclude that sclerotome-derived VSMC progenitors contribute to the embryonic HSC specification niche, most likely by providing unknown HSC inductive signals.

## Main

Haematopoietic stem cells (HSCs) are progenitor cells that both self-renew and are capable of repopulating all haematopoietic lineages in vertebrates^1–3^. HSCs form the basis of transplant therapies for haematologic diseases including leukaemia, haemoglobinopathies, congenital neutropenias^4,5^, and sickle cell disease^6^. Successful directed differentiation of pluripotent stem cells, such as induced pluripotent stem cells (iPSCs), to an HSC fate could provide an unlimited source of HSCs for autologous and allogeneic transplants, drug studies, and disease modeling^7^. The recent advent of multiple promising or successful gene therapy trials also points to HSCs as a desirable vector for delivery of genetically edited and permanent haematopoietic reconstitution^8–10^. Moreover, proliferative populations, such as cells undergoing directed differentiation, are more receptive to genetic correction or modification. An intuitive means of directing HSC differentiation *in vitro* is to attempt to recapitulate normal specification during embryonic development, and intense research has focused on understanding this process^1,7^.

Across vertebrate phyla including human, mouse, and zebrafish, HSCs are specified from specialized endothelium known as haemogenic endothelium, found in the floor of the primitive descending aorta in mammals, or its cognate vessel, the dorsal aorta (DA) in zebrafish^1^. Haemogenic endothelium derives from splanchnopleural mesoderm in mammals and lateral plate mesoderm in anamniotic vertebrates, and its competence is primed by earlier embryonic signaling that leads to a primitive DA capable of sensing and responding to requisite inductive signals, provided by a localized HSC specification niche, which is distinct in location and purpose from the adult HSC homeostasis niche found in the mammalian bone marrow or zebrafish kidney marrow^1^. The signaling events supporting haematopoietic specification, insofar as they are currently understood, are strongly conserved from fish to mammals^1,7^. We previously showed that neural crest cells contribute to the HSC specification niche, and that treatments that prevent crest proximity to the haemogenic endothelium result in diminished or abrogated specification of HSC precursors, identifying them as a critical component of this embryonic specification niche^11^.

Although many essential endogenous signaling pathways have been identified that contribute to HSC specification—including Notch, bone morphogenetic protein (Bmp), sonic hedgehog (Shh)-vascular endothelial growth factor A (Vegfa), nitric oxide (NO), canonical wingless-type MMTV integration (Wnt) signaling, transforming growth factor β (Tgfβ), and inflammatory signaling—successful high efficiency *in vitro* directed differentiation of normal HSCs by treatment of iPSCs or *in vitro*-generated haemogenic endothelium with exogenous factors (rather than permanent genetic modification by viral transduction with transcription factors) has yet to be reported, suggesting that critical signals remain to be identified^1,7,12–15^.

Non-canonical Wnt signaling also regulates HSC emergence in zebrafish. We previously showed that HSC specification depends on Wnt16^16^. Although Wnt16 morphants appear grossly normal and retain formation of nearly all developing tissues, including the DA, pronephros, and myotome, the most ventral compartment of the somite – the sclerotome – is severely disrupted. These results are consistent with the possibility that proper sclerotome specification or morphogenesis is required for HSC specification, most obviously by contributing to the HSC specification niche and providing HSC-inductive signals to the haemogenic endothelium.

The sclerotome comprises the ventral portion of vertebrate somites, metameric blocks of mesoderm in the trunk of the developing embryo, and it houses progenitors for multiple adult tissues including vertebral column, meninges, tendons, intervertebral discs, ribs, and sternum^17–19^. Association of sclerotome cells with the descending aorta was first observed just over 150 years ago^20^, and sclerotome has been proposed to give rise to VSMCs in chick and mouse within the last decade, but these studies relied upon heterotypic grafts and observation of lineage markers in fixed embryos^21–24^. More recently, work in zebrafish identified trunk VSMCs of the DA as being sclerotomally derived, using a transgenic putative sclerotome reporter developed with Medaka regulatory elements from the orthologous *Twist* gene^25^. Although these studies support the assignment of VSMCs of the DA as sclerotomal in origin, they do not address the kinetics of the initial association between VSMC precursors and the primitive DA, which is a critical consideration for the evaluation of VSMC precursors as HSC specification niche cells.

Here we have generated a novel transgenic sclerotome reporter using regulatory elements derived from the zebrafish orthologue of the paradigmatic sclerotome marker, *Pax1*. Our results confirm the sclerotomal origin of DA VSMCs and clearly define the timing of initial contact between the DA and VSMC precursors. In functional studies, we have moreover disrupted sclerotome patterning to observe the consequences for definitive haematopoietic programming. We show that knockdown of specific sclerotome patterning genes leads to both loss of specification of HSC precursors in the haemogenic endothelium, and later loss of VSMCs of the DA. Our results support a model in which sclerotome-derived VSMC precursors contribute to the embryonic cellular HSC specification niche and present critical inductive signals.

## RESULTS

### Sclerotome-derived cells associate with the DA at the right time and place to contribute to the HSC specification niche

To better understand the dynamic localization and tissue potential of sclerotome cells, we generated a transgenic zebrafish line (Fig. 1, Supplementary Fig. 1) in which expression of the Neon fluorophore is driven by regulatory elements derived from the prototypical sclerotome marker gene *pax1a* (*Tg(en.pf1-2.5pax1a:Neon)^sj6^*, hereafter referred to as *pax1a:Neon*; Supplementary Fig. 1a-c). The mouse *Pax1* gene is one of the oldest and best characterized sclerotome markers^17,18,26,27^. *Pax1* genes are duplicated in zebrafish, and of the two paralogous zebrafish *pax1* genes, only *pax1a* exhibits expression in the sclerotome^16,28^. We were able to identify an 891 bp enhancer (*pf1*), located approximately 138.5 kilobasepairs 3’ to the annotated *pax1a* coding locus, based on comparative genomics and published studies of *Pax1* regulation (Supplementary Fig. 1a)^29^. When combined with a 1.9 kilobase (kb) region located directly 5’ to the *pax1a* coding region 5’-most initiator, along with exon 1 and intron 1 of the endogenously occurring *pax1a* gene, which displays alternative initiation exons (Supplementary Fig. 1a-c), we were able to direct robust sclerotome-specific expression of the *Neon* fluorophore transgene, which additionally lacked the unwanted pharyngeal endoderm expression characteristic of both *pax1a* and the paralogous gene *pax1b* (Fig. 1a-d; Supplementary Fig. 1d-k). The *pax1a:Neon* transgenic provides a tool for tracking sclerotome morphogenesis in real time in living embryos, and has a more robust, consistent, and defined pattern than the *Ola.Twist:GAL4* reporter previously used^25,30^.

**Figure 1.**
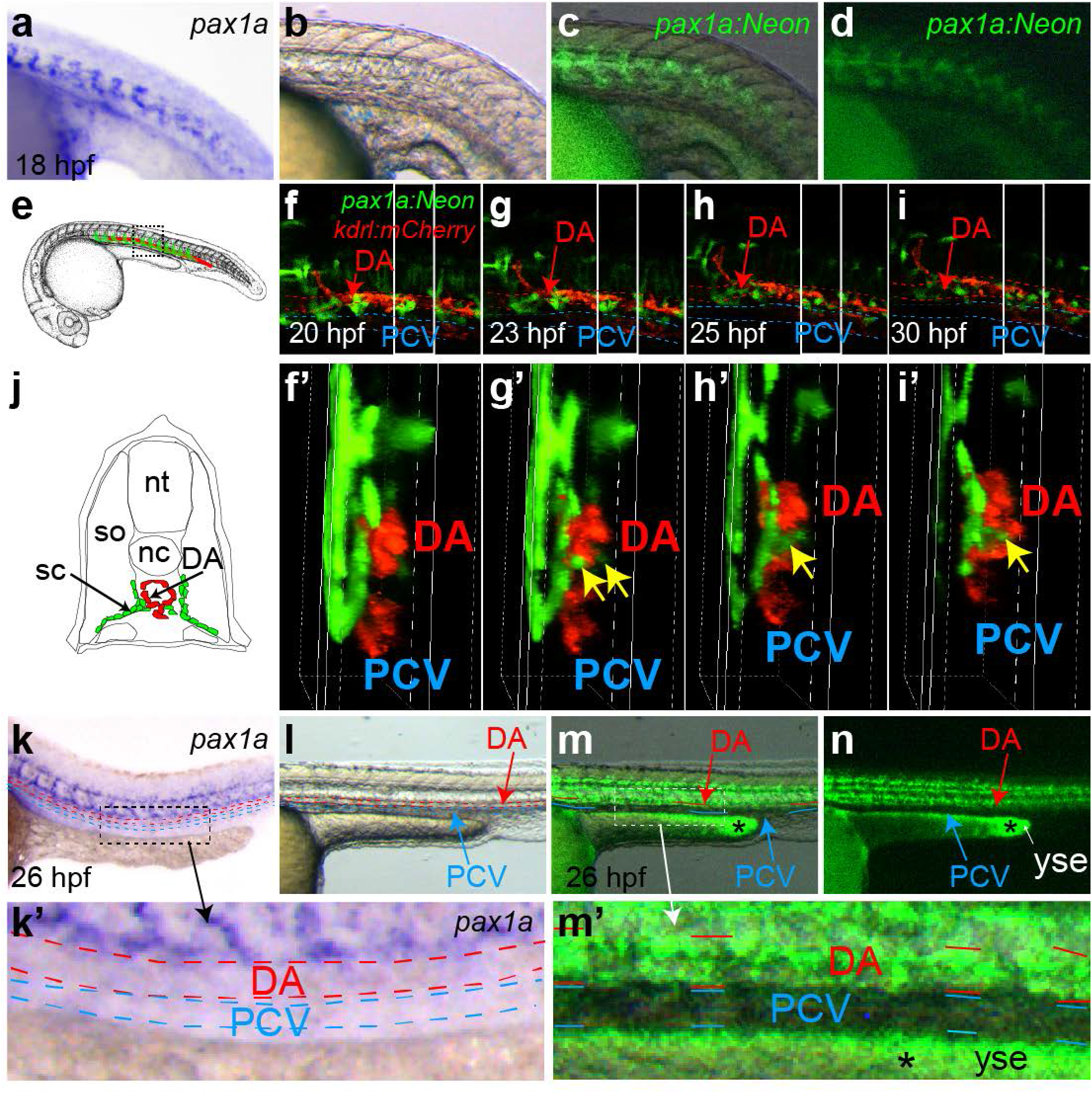
A *pax1a:Neon* transgenic reports sclerotome morphogenesis. Comparison of endogenous trunk expression of the *pax1a* transcript by whole mount *in situ* hybridisation (a) at 18 hpf with with brightfield (b), Neon-brightfield dual channel (c), and Neon fluorescence (d) in *pax1a:Neon* transgenics. Schematics (e, j) and sequential stills (f-i’) of time lapse confocal microscopy of a double *pax1a:Neon; kdrl:mCherry* transgenic from 20 - 30 hpf. Lateral schematic (e) illustrates the trunk region depicted in f-i, while a transverse schematic (j) illustrates the virtual transverse sections depicted in f’-i’ virtually projected from the region of the grey boxes in f-i. Green sclerotome-derived cells contact the ventral side of the DA (yellow arrows) beginning at 23 hpf (g’). At 26 hpf and beyond, the *pax1a* transcript is no longer detectable in cells contacting the DA (k, k’, l), but the Neon fluorophore persists (m, m’, n). DA, dorsal aorta (red arrows f-i, l-n); PCV, posterior cardinal vein (blue arrows, l-n); nc, notochord; nt, neural tube; sc, sclerotome; so, somite; yse, yolk sac extension; * autofluorescence in yse. Lateral images, dorsal up, anterior left (a-i; k-m’), transverse images, dorsal up (f’-i’; j). 80X (a-d; k-n). 100X (f-i). 160X (f’-i’). Digital zoom of boxed regions in k and m (k’, m’).

Using the *pax1a:Neon* sclerotome reporter animals, we were able to track the exact progress of sclerotome migration beginning from late somitogenesis through and beyond 24 hours post fertilization (hpf), the time window relevant to the establishment of the cellular HSC specification niche. Sclerotome cells labeled by Neon began associating with the DA by around 22 hpf, where they initiated and maintained contact with the ventral aspect of the DA (Figure 1e-j; Supplementary Movies 1-4). Although the *pax1a* transcript appears to be actively degraded in the DA-associated cells, the Neon fluorophore perdures, revealing specific association of sclerotome-derived cells with the DA and not the posterior cardinal vein (PCV) beneath (Fig. 1k-m’). As HSC precursors are found predominantly in the ventral portion of the DA, the preferential association of sclerotome cells with this vessel suggests the possibility of specific signaling contact with the HSC progenitor population in the haemogenic endothelium.

### Sclerotome cells become VSMCs of the DA

Previous studies indicated that VSMCs may be sclerotomally derived^20–25^. To explicitly examine the VSMC fate potential of DA-associated Neon^+^ cells in our *pax1a:Neon* animals, we crossed them to a VSMC *acta2:mCherry* reporter line^31^. Confocal imaging of double transgenic animals confirmed that Neon^+^ sclerotome cells, which were already associated with the DA by 22 hpf, began to express mCherry, indicating their overt differentiation to VSMCs, by 3 days post fertilization (dpf), and were easily observable by 4 dpf (Fig. 2a-f) and beyond through at least 8 dpf (not shown) both by confocal microscopy and as clear double positive flow cytometric populations at both times (Fig. 2g). These results indicate that sclerotome-derived cells associate with the DA at 24 hpf, and contribute to arterial VSMCs, preferentially localized along the ventral surface of the DA at 3 dpf and beyond, confirming and extending previous studies. This population of VSMC progenitors is thus well positioned to provide HSC inductive signals and contribute to the HSC specification niche. Interestingly, overt differentiation of sclerotome cells to VSMCs in zebrafish at around 3.5 days post fertilization (dpf), as judged by initiation of expression of the mCherry reporter and morphological appearance of VSMCs^25,32,33^, appears to coincide with the approximate time at which new HSC precursors cease to be specified in and emerge from the DA haemogenic endothelium.

**Figure 2.**
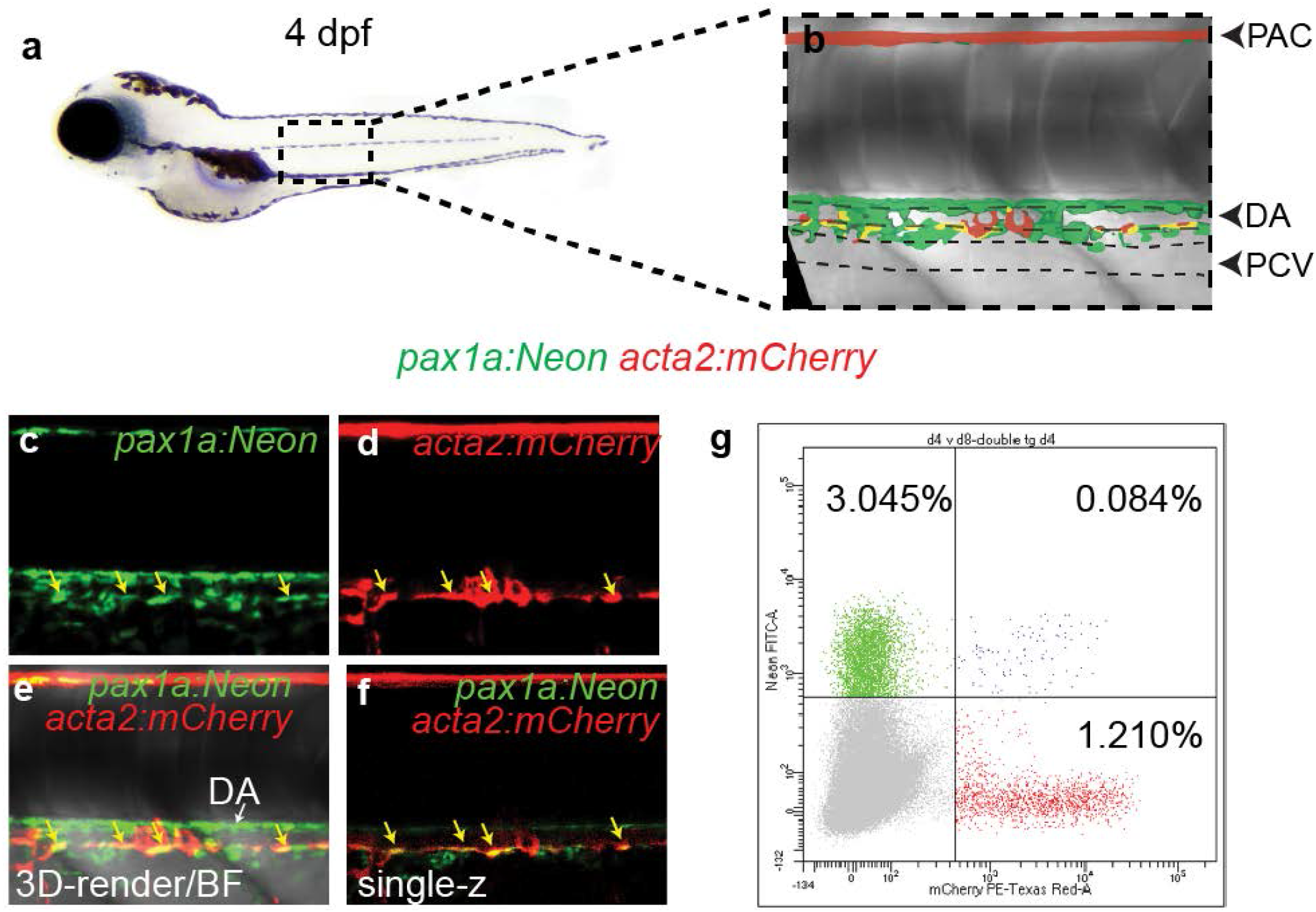
Sclerotome contributes to vascular smooth muscle cells. Schematic representation (a, b) of the trunk region of a 4 dpf *pax1a:Neon*; *acta2:mCherry* embryo. The regions of the DA, PCV, and parachordal vessel (PAC) are indicated (b). Confocal images (c-f) reveal Neon^+^ mCherry^+^ double positive cells in a 0.2 µm single-z, demonstrating that sclerotome-cells have initiated expression of the smooth muscle marker *acta2*. Flow cytometry analysis of the trunk region of double transgenic animals confirms the double positive population 4 dpf. Neon (c) and mCherry (d) max projections, three-channel 3D-render (e), and two-channel 0.2 µm single-z (f) images. 160X lateral images, dorsal up, anterior left (b-f). DA, dorsal aorta; PCV, posterior cardinal vein; PAC, parachordal vessel.

### Morpholino knockdown of sclerotome genes disrupts sclerotome patterning

To understand the fate potential of sclerotome and its inductive signaling potential, we wanted to establish a set of tools to disrupt normal sclerotome specification and morphogenesis. We determined a number of sclerotome-specific genes with likely requirement in sclerotome patterning, including *pax1a*, *pax9*, *twist1b*, *twist2*, *foxc1a*, *foxc1b*, and *snai2*. In choosing genes for sclerotome perturbation, we wanted to exclude genes also expressed in the endothelium, because they might affect haematopoietic potential cell-autonomously. Although all genes examined appeared not to be expressed in the endothelium based on whole mount *in situ* hybridisation at 24 hpf, we discovered that a subset of genes (*foxc1a*, *foxc1b*, and *snai2*) showed plausible expression by reverse transcriptase polymerase chain reaction (RT-PCR) analysis of highly purified trunk endothelial cells isolated by flow cytometry from *kdrl:mCherry; fli1a:EGFP* double transgenic zebrafish (Supplementary Fig. 2). The mCherry EGFP (*kdrl*, *fli1a*) double positive population was particularly of interest, because it alone expresses consistent levels of the pre-HSC markers *gata2b* and *runx1*, suggesting that HSC precursors might be preferentially located within this fraction. As *pax1a*, *pax9*, *twist1b*, and *twist2* were not found within the endothelial compartment based on *in situ* or flow cytometric RT-PCR analysis, we limited ourselves to knockdown of these genes in an attempt to specifically disrupt sclerotome patterning.

To disrupt sclerotome formation we took advantage of established antisense morpholino oligonucleotides^34,35^ targeting *pax9* (Pax9MO), *twist1b* (Twist1bMO), and *twist2* (Twist2MO), as well as a newly designed morpholino that prevented transcript maturation of *pax1a* (Pax1aMO; Supplementary Fig. 3), all of which we suspected might disrupt sclerotome patterning. Published studies have raised concerns about potential “off-target” effects of morpholinos^36,37^, and an established set of criteria have been developed to provide guidance for when to draw conclusions about *bona fide in vivo* genetic loss-of-function effects attributable to genes based on morpholino knock down approaches^38^. Although we have obtained mutants for most of the genes in our set and performed specificity studies (see below), it is important to emphasize that in this case we only wished to find tools suitable for preventing normal association between sclerotome and haemogenic endothelium of the DA, irrespective of whether the morpholinos provide a means to draw conclusions about the role of the specific genes targeted. Thus, whether or not our morpholinos reveal the actual *in vivo* functions of the targeted genes, they are useful if they cause robust and reproducible disruptions in sclerotome patterning allowing us to prevent association of sclerotome with the haemogenic endothelium.

We analysed sclerotome integrity in animals injected with Pax1aMO, Pax9MO, Twist1bMO, and Twist2MO in comparison to wild type, uninjected embryos at 22 hpf (Fig. 3; Supplementary Table 1). Injection of each morpholino was sufficient to severely perturb normal sclerotome specification and/or morphogenesis. We cannot rigorously distinguish failure of sclerotome specification from migration, but it appears that both are likely in play. This panel of morpholinos therefore provides the tools to test the effect of defects in sclerotome patterning on both cell-autonomous tissue contribution of sclerotome to VSMCs, and non-cell-autonomous induction of haematopoietic commitment in the haemogenic endothelium by acting as HSC specification niche cells.

**Figure 3.**
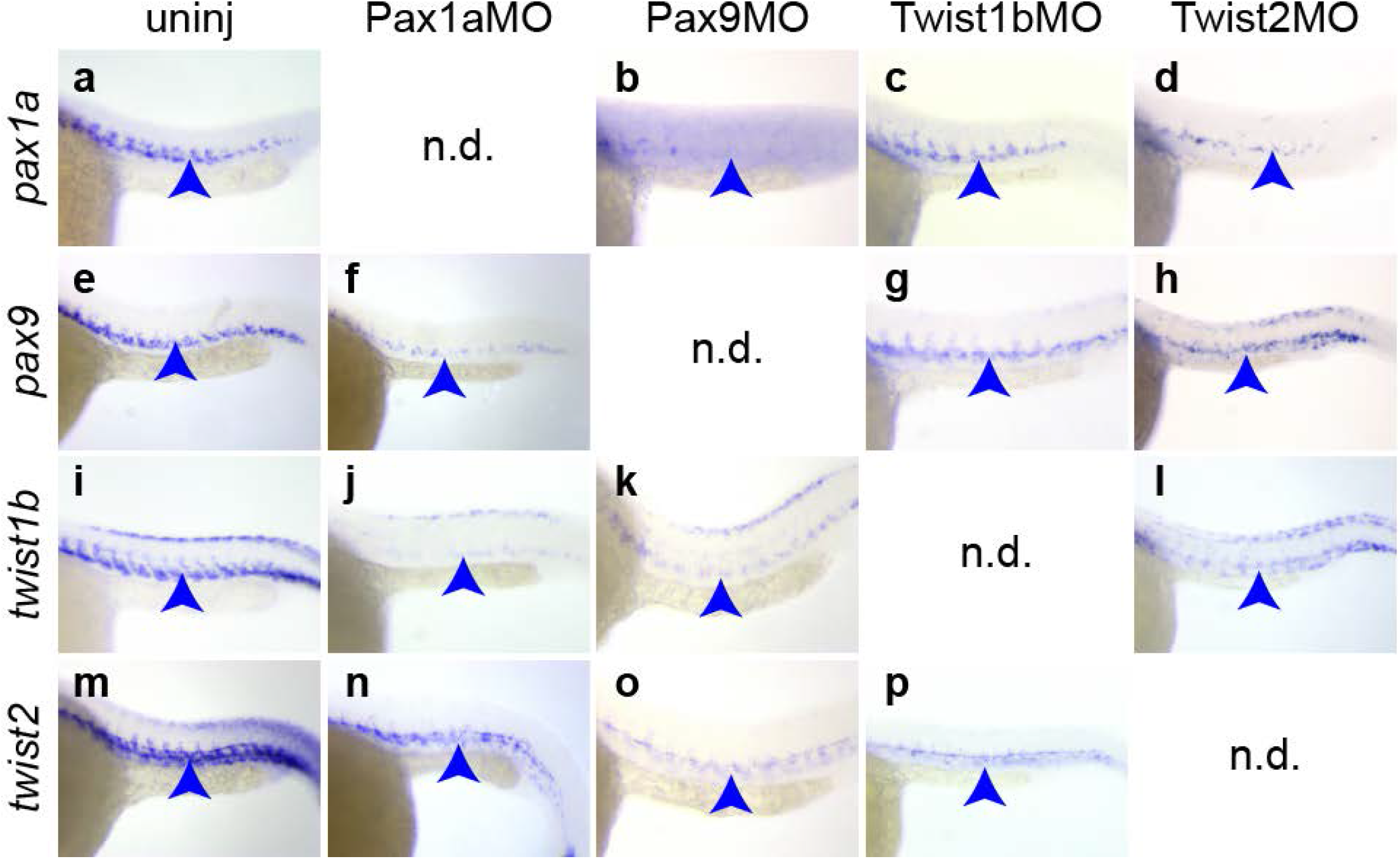
Morpholinos targeting sclerotome genes disrupt sclerotome patterning. Sclerotome marker expression (*pax1a*, a-d; *pax9*, e-h; *twist1b*, i-l; *twist2*, m-p) in embryos either uninjected (a, e, i, m), or injected with Pax1aMO (f, j, n), Pax9MO (b, k, o), Twist1bMO (c, g, p) or Twist2MO (d, h, l). Sclerotome gene expression in morpholino-injected embryos is consistently disrupted in comparison to uninjected embryos. The regions of presumptive sclerotome expression are indicated (blue arrowheads). n.d., not done. 80X lateral images, dorsal up, anterior left.

### Normal sclerotome patterning is required for VSMC investment of the DA

Based on our observation that sclerotomal cells contribute to VSMCs of the DA, we predicted that we might see defects in VSMC development in embryos injected with our morpholinos disrupting sclerotome patterning. To test this possibility, we injected double transgenic *fli1a:EGFP;acta2:mCherry* animals^31^ (Fig. 4a,b) with Pax1aMO, Pax9MO, Twist1bMO, and Twist2MO to be able to observe both endothelium and VSMC development in animals where sclerotome patterning is disrupted (Fig. 3). By 5 dpf we could easily see the appearance of mCherry^+^ VSMCs along the ventral floor of the DA in wild type, uninjected animals (Fig. 4a-f). In contrast, although endothelium of the DA and other major vessels was relatively normal in morphant animals, few or no VSMCs could be observed associated with the DA (Fig. 4g-w). Interestingly, VSMCs associated with the gut and parachordal vessels appeared relatively intact, suggesting that these cells are not sclerotome-derived. Our results indicate that sclerotome contributes specifically to VSMCs of the DA.

**Figure 4.**
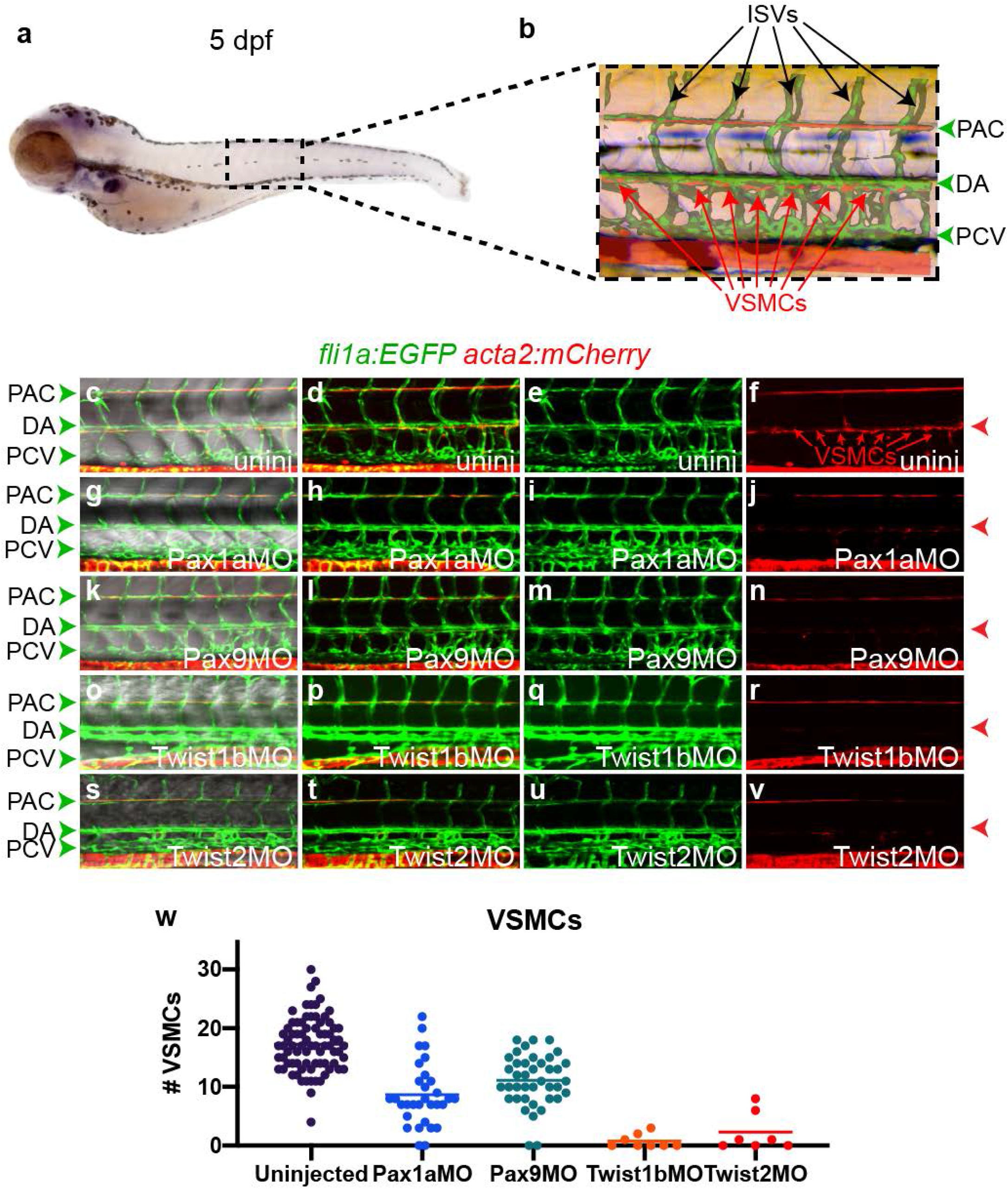
Loss of VSMCs of the DA in embryos with sclerotome disruption. Schematic of the trunk regions and vascular anatomy in a 5 dpf embryo (a, b). VSMCs of the DA are present at 5 dpf in uninjected (c-f), but absent in embryos injected with Pax1aMO (g-j), Pax9MO (k-n), Twist1bMO (o-r), or Twist2MO (s-v). The level of the DA is indicated at right (red arrowhead, c-v). A plot depicting the number of VSMCs counted in embryos for each condition is presented with the mean indicated (w). 160X lateral images, dorsal up, anterior left (b-v). DA, dorsal aorta; PCV, posterior cardinal vein; ISVs, intersegmental vessels (black arrows, b); PAC, parachordal vessel; VSMCs, vascular smooth muscles (red arrows, b, f).

### Normal haemogenic endothelium in sclerotome morphants

Our goal was to determine whether sclerotome might contribute to the HSC specification niche, so we wanted to understand whether normal haemogenic endothelium, capable of responding to niche signaling was present in morphants. We therefore examined the state of gross embryonic patterning with special emphasis on haemogenic endothelium. Myotome formation (*myod1*; Fig. 5a-e; Supplementary Table 2) and primitive erythropoiesis (*gata1a*; Fig. 5f-j; Supplementary Table 2) were both intact and normal in morphant compared to uninjected, wild type embryos. Similarly, endothelium (Fig. 4c-e,g-i,k-m,o-q,s-u; *cdh5* Fig. 5k-o; Supplementary Table 2) with arterial identity (*efnb2a* Fig. 5p-t; Supplementary Table 2) and competent to respond to key instructive Notch signals (*notch1b* Fig. 5u-y; Supplementary Table 2) was also present. Sclerotome morphants exhibited normal early circulation of primitive erythrocytes (not shown). Thus, normal haemogenic endothelium is present in sclerotome morphant animals at the time of definitive haematopoietic initiation.

**Figure 5.**
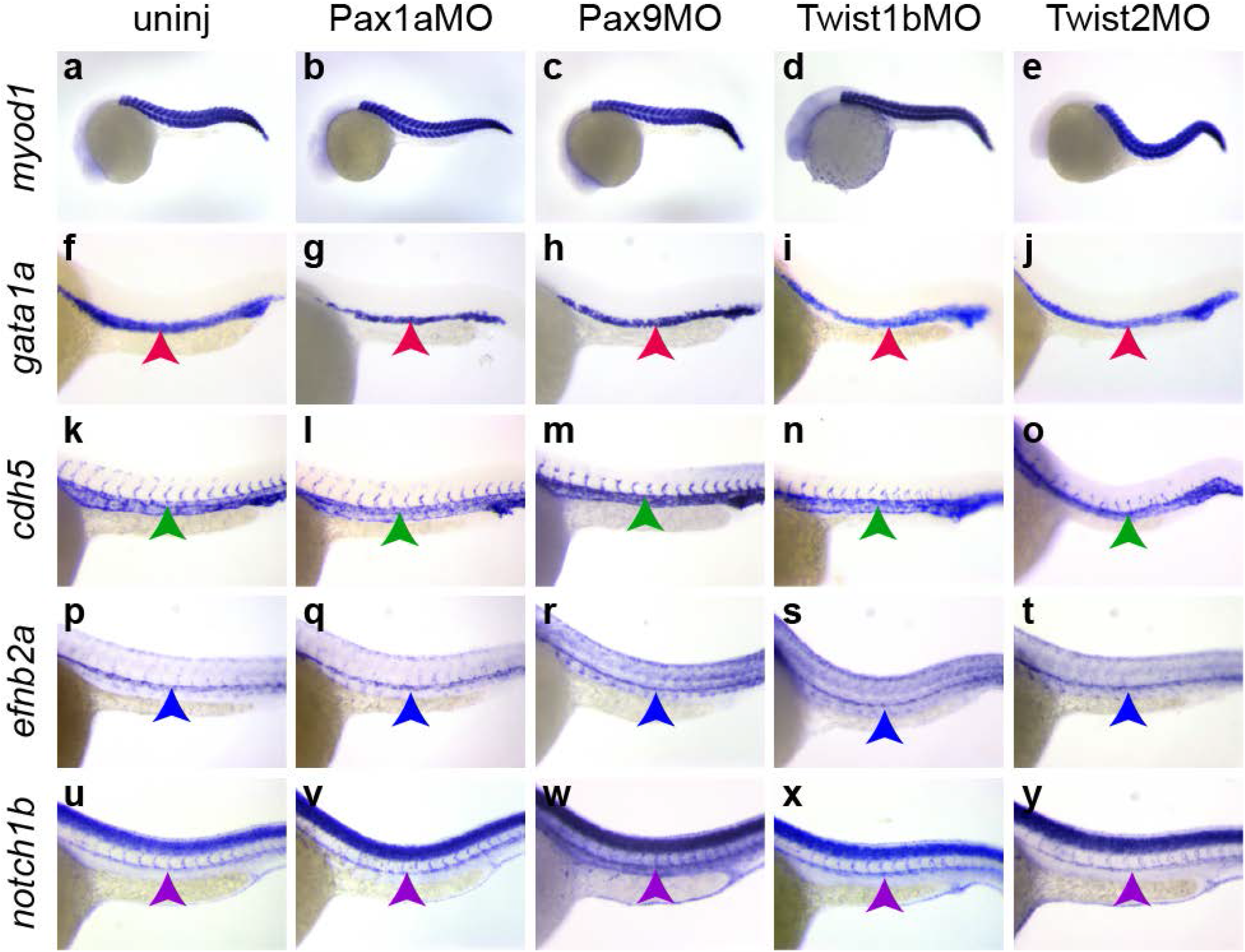
Normal gross patterning of sclerotome-disrupted embryos. Examination of the state of myotome (*myod1*, a-e), primitive erythropoiesis (*gata1a*, red arrowheads, f-j), endothelium (*cdh5*, green arrowheads, k-o), arterial specification (*efnb2a*, *notch1b*, blue and purple arrowheads, p-y), and Notch signaling capability (*notch1b*, purple arrowheads, u-y), in uninjected or embryos injected with the morpholinos indicated above each column, reveals normal myotome and primitive erythropoiesis in all individuals, as well as intact, Notch-signaling competent, arterial endothelium. Lateral images, dorsal up, anterior left. 40X (a-e) or 80X (f-y).

We previously showed that neural crest cells contribute to the HSC specification niche^11^. We therefore wished to examine the status of neural crest patterning in morphants. We injected embryos with Pax1aMO and Pax9MO and examined neural crest patterning by whole mount *in situ* for the neural crest marker *crestin*. For both morpholinos, neural crest patterning was normal in morpholino-injected animals compared to uninjected controls (Supplementary Fig. 4; Supplementary Table 3). As *twist1b* and *twist2* are expressed in neural crest populations, it is likely that neural crest migration is affected in animals with these genes knocked down, however, the presence of normal neural crest patterning in Pax1aMO and Pax9MO animals allowed us to determine haematopoietic development defects uniquely due to disruption of sclerotome patterning.

### Sclerotome cells contribute to the HSC specification niche

We hypothesized that sclerotome patterning is required for specification of HSCs by contributing VSMC precursors to the HSC specification niche; we therefore wanted to determine whether any problems in normal definitive haematopoietic development could be observed in animals with sclerotome defects, notwithstanding the apparently normal state of primitive haematopoiesis (Fig. 5f-j; Supplementary Table 2). HSC precursors in the haemogenic endothelium can be recognized by expression of *runx1*, which has been shown to be required for the initiation of definitive haematopoietic programming and normal HSC specification across vertebrate phyla^1^. In comparison to wild type, uninjected animals, *runx1* expression was very significantly reduced in all sclerotome morphants at 24 hpf (Fig. 6a-e; Supplementary Tables 4, 5). These results indicate that proper sclerotome patterning is required for the emergence of *runx1*^+^ HSC precursors in the haemogenic endothelium.

**Figure 6.**
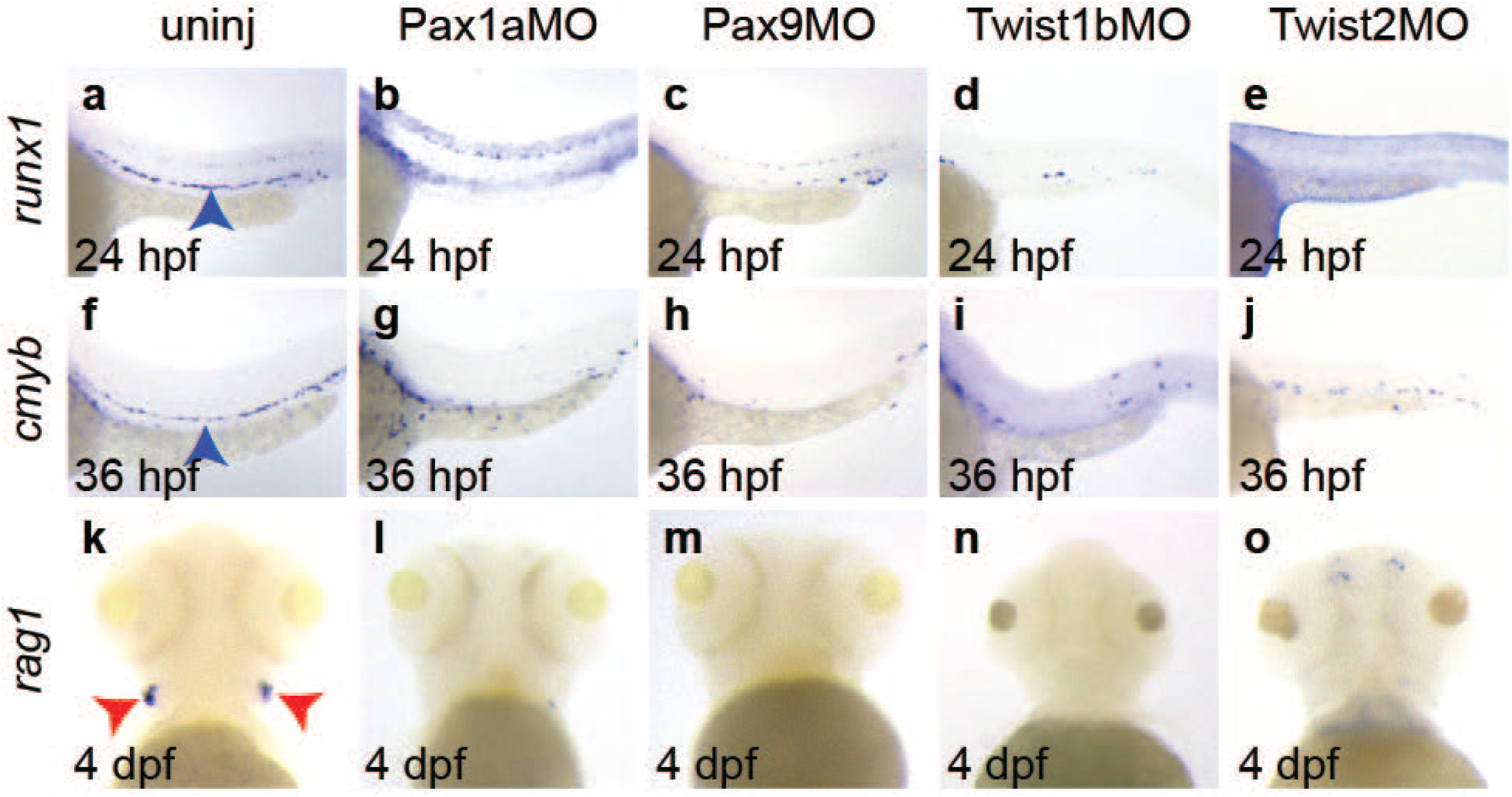
HSC specification requires sclerotome. Examination of the state of definitive haematopoiesis by examination of the HSC precursor marker, *runx1* (a-e, blue arrowhead) at 24 hpf, emerging HSC marker, *cmyb* (f-j, blue arrowhead) in the trunk DA at 36 hpf, and T lymphocyte marker, *rag1* (k-o) in the thymus (red arrowheads) at 4 dpf reveal loss of definitive HSC precursors and T lymphocytes in sclerotome-disrupted embryos. a-j, 80X lateral images, dorsal up, anterior left. k-o, 100X ventral images, anterior up.

We also examined *runx1* expression in animals carrying predicted loss-of-function mutations in a subset of the sclerotome genes we targeted by morpholino, including newly generated *pax1a* (Supplementary Fig. 5) and *twist2* (Supplementary Fig. 6) mutants, as well as an existing *twist1b* mutant^39^, but in contrast to morpholino knockdown, we saw no apparent defects in *runx1* expression in homozygous mutants (Supplementary Fig. 7a,b,e,f,i,j; Supplementary Table 5). Differences in phenotype between mutants and morpholino-injected animals could be due either to off-target morpholino effects, acute genetic compensation termed “transcriptional adaptation”^40,41^, or other forms of compensation in the mutants^42^. To distinguish between these possibilities, we injected wild type, heterozygous, and homozygous mutant embryos with morpholinos targeting the mutant gene, in accordance with published guidelines for determining specificity of morpholino effects^38^. We examined *runx1* expression in uninjected animals, and scored expression (blind) followed by individual genotyping. Although morpholinos caused robust decrease of *runx1*^+^ precursors in wild-type or heterozygous animals, they were no longer effective in homozygous mutants (Supplementary Fig. 7a-l; Supplementary Table 5), indicating the presence of compensatory gene expression, or other modes of compensation. Our results strongly suggest that failure to recapitulate the morpholino loss-of-function phenotype in the *pax1a*, *twist1b*, and *twist2* mutants is the consequence of compensation. Irrespective of compensation, morpholino injection readily caused the main effect we wished to test – disruption of normal sclerotome patterning – and thus provided a means to test the necessity of that patterning for HSC specification.

To verify compromised definitive haematopoiesis in sclerotome morphants, we examined a second marker of HSC emergence, *cmyb*, and found it also to be severely reduced (Fig. 6f-j; Supplementary Table 4). We also examined the emergence of T lymphocytes by expression of *rag1* in the bilateral thymi of 4 dpf zebrafish. Like HSC precursor markers, *rag1* was greatly diminished or absent in sclerotome morphants (Fig. 6k-o; Supplementary Table 4), confirming that the emergence of *runx1*^+^ HSC precursors and other definitive haematopoietic progenitors with lymphoid potential had been severely decreased. We conclude that proper sclerotome patterning is required for induction of definitive haematopoietic potential from existing arterial haemogenic endothelium.

## DISCUSSION

Our previous studies demonstrated that defects in sclerotome patterning are coincident with loss of HSCs^16,43^ pointing to the hypothesis that sclerotome specification or morphogenesis might somehow be required for HSC specification. We generated transgenic animals to precisely map the timing and location of sclerotome tissue morphogenesis in explicit comparison to emergence of HSC precursors in the haemogenic endothelium of the primitive DA. Our results demonstrate by live imaging that individual sclerotome precursors originating in the ventral somite migrate dorsally, with some inserting themselves between the DA and the PCV and making contact with ventral DA endothelium by 23 hpf, the time of onset of definitive haematopoiesis. Other sclerotome-derived cells continue dorsalward, presumably to contribute to other adult sclerotome-derived tissues including vertebral column and somite connective tissue. Timing and location of sclerotome contact with the DA thus positions this population as candidates to present haematopoietic instructive signals to HSC precursors of the haemogenic endothelium.

Our studies show that cells in contact with the DA form or contribute to VSMCs of the DA, in confirmation of previous work^21–25,32^. When sclerotome patterning is disrupted, VSMCs do not invest the DA, although other vessels such as the parachordal are unaffected, suggesting a different origin for those VSMCs. Surprisingly, VSMC-fated precursors appear to reach their anatomical site of differentiation, in contact with the DA, by 23 hpf, far earlier than the time when they overtly differentiate to VSMCs as judged by morphology and expression of VSMC markers^25,32,33^. Conceivably, signaling from the DA (and other local cell types) to the VSMC precursors instructs this differentiation process. As emergence of *runx1*^+^ precursors from the haemogenic endothelium appears to terminate around 3.5 dpf, approximately the same time as VSMC differentiation, it is possible that VSMC precursors – and not differentiated VSMCs – are uniquely competent to present instructive signals.

In addition to VSMCs, sclerotome contributes to diverse adult tissues, including vertebral column, meninges, tendons, intervertebral discs, ribs, and sternum^17–19^. The timing of when fate potential within the sclerotome is established is not well understood, although very early sclerotome exhibits differences in migratory behaviour and overlapping but distinct compartmental gene signatures, suggesting that sclerotome fates might be established or in progress earlier than previously supposed. In the future, it will be important to address whether particular combinations of genes cooperate to drive discrete lineage commitments within the sclerotome, and determine how early those fates are identifiable.

Our results demonstrate that one set of genes, including *pax1a*, *pax9*, *twist1b*, and *twist2*, appears to drive commitment to VSMCs, and that sclerotome patterning defects resulting from knockdown of these genes leads to both loss of VSMCs, and earlier loss of HSC precursors. A simple model explaining these results is that sclerotome-derived VMSC precursors migrate to and contact the DA to instruct specification of the earliest HSC precursors from the haemogenic endothelium, thereby acting as an embryonic HSC specification niche.

We recently described another cell type that contributes to the embryonic HSC specification niche, the neural crest^11^. Perhaps unsurprisingly, as both cell types are highly migratory and exhibit diverse fate potentials, there is significant overlap in gene expression between sclerotome and neural crest, including expression of genes that regulate epithelial-to-mesenchymal transition, such as *twist1b*, *twist2*, *snai1b*, and *snai2* For these genes, we expect that loss-of-function likely impairs HSC specification by preventing migration of both neural crest and sclerotome. Despite significant overlap in genes that regulate sclerotome and neural crest, we are also able to isolate and distinguish unique requirements for both cell types through knockdown of genes with specific expression and tissue-patterning roles, for example, *pax1a* and *pax9*. Together, our results indicate that the embryonic specification niche requires cooperative contribution of multiple distinct cell types, each necessary but not sufficient.

How do sclerotome and other niche cells cooperatively instruct HSC specification? The simplest model is that distinct cell populations each present one or more short-range signals that are combinatorially required to initiate haematopoietic programming. However, more complicated models are conceivable. Perhaps sclerotome-derived VSMC precursors direct neural crest maturation that potentiates expression of HSC specification signals. Or perhaps sclerotome or neural crest cells modify the local environment to potentiate signaling or other requirements for haematopoietic programming. Future studies will examine the exact nature of interactions between sclerotome, neural crest, and the DA to determine how each cell type influences the HSC specification environment and programming of cell fates.

## METHODS

### Zebrafish husbandry

Wild type (AB, WIK, TL), *Tg(en.pf1-2.5pax1a:Neon)^sj6^*, *Tg(acta2:mCherry)^uto5^*, *Tg(fli1a:EGFP)^y1^*, *Tg(−6.5kdrl:mCherry)^ci5^*, *pax1a^sj7^*, *twist1b^el570^*, and *twist2^sj8^* lines were maintained as previously described^44^, under an institutionally approved (IACUC) animal use protocol (Protocol 555), and in compliance with all relevant AVMA guidelines. Embryos were obtained by natural and timed spawnings and raised as previously described^44^. Embryos were not preferentially selected by sex.

### Transgenic and mutant line generation

The *Tg(en.pf1-2.5pax1a:Neon)^sj6^* line was generated by Tol2 transposon-mediated transgenesis^45^ using a construct containing Tol2 cis arms flanking the 891 bp putative *pf1* enhancer element located at Ch17:42416407-42417279 (Ensembl GRCz11), upstream of a minimal promoter fragment from the endogenous *pax1a* locus spanning the region from −2456 to +27 relative to the ATG of the *pax1a-201* splice variant, thus containing both potential initiation codons and intron 1 of the *pax1a-202* splice variant, followed by the sequence encoding the mNeon fluorophore ^46^, and a heterologous *SV40 polyA* signal motif (Supplementary Fig. 1). Transgenesis constructs (25 pg) were co-injected with Tol2 mRNA (25 pg) into TL embryos and transgenic animals were identified by sclerotome fluorescence and bred to single insertions.

The *pax1a^sj7^* (Supplementary Fig. 5) and *twist2^sj8^* (Supplementary Fig. 6) mutant lines were made by CRISPR/Cas9-mediated deletion of locus-specific fragments, including a 2614 bp fragment from the 5’-UTR through exon 4 of the *pax1a-202* splice form (exon 3 of the *pax1a-201* splice form; GRCz11 17:42274369-42276984) and two fragments of *twist2*, a 21 bp fragment from +33 to +53 and a 372 bp fragment from +87 to +458 relative to the ATG initiator (GRCz11 9:45839482-45839502 and 9:45839536-45839907), following injection of *in vitro* transcribed gRNAs targeting the *pax1a* sequences 5’-CTCCAGCAAACATCACCCTGCGG-3’ and 5’-AGTCTTACCCACCCTGGCGGTGG-3’ or *twist2* guide sequences 5’-GACAGCAGAAAAGGTTCGGGAGG-3’ and 5’-GTTACGCGTTTTCAGTGTGGAGG-3’ as previously described^47^. *In vitro* synthesized sgRNAs were transcribed using T7 polymerase, and 200 pg of each was co-injected with 500 pg Cas9 protein (IDT) into AB (*pax1a*) or WIK (*twist2*) embryos. Stable mutants were identified and validated by PCR and sequencing of the loci. *Pax1a* stable mutants were genotyped using multiplexed PCR on genomic DNA from individual or pooled embryos, or fin-clipped adults, using the F3b (5’-CCCATTGCTCTCCGCATAAC-3’), F3c (5’-GCCAAGCACGGATAATGTGT-3’), and R3 (5’-AGGCCACTGTTGAGATCATGC-3’) primers (Supplementary Fig. 5; Supplementary Table 6) to amplify a 511 bp (wild-type, F3c+R3) or 322 bp (mutant, F3b+R3) band respectively. *Twist2* mutants were genotyped using the primers, twist2 F 5’-ATGGAAGAGAGTTCTAGCTCTCCC-3’ and twist2 R 5’-CTAGTGGGACGCAGACATCG-3’ to amplify a 492 bp wild type or 96 bp mutant band (Supplementary Fig. 6; Supplementary Table 6).

### In situ probes and whole mount *in situ* hybridisation

Whole mount *in situ* hybridisation was performed as described previously^16^. Briefly, digoxigenin and fluorescein labeled ribo-probes were transcribed from 5’-linearised probe constructs using T3, T7, or SP6 RNA polymerase. Embryos were fixed, permeabilised, hybridised with probe, and developed. The *twist1b*, *twist2*, *myod1*, *cdh5*, *efnb2a*, *notch1b*, *runx1*, *cmyb*, *rag1*, and *gata1a* probes were described previously^16^. *Pax9* was cloned from 24 hpf cDNA using the primers pax9F 5’-TCTAGAATGGAGCCAGCCTTT-3’ and pax9R 5’-ATGGATCCTCATAGAGCTGAAGCCACCAG-3’ (Supplementary Table 6) and cloned by TOPO-TA to the pCRII vector (Invitrogen) to create pCRII *pax9*. The *mNeon* probe was made by amplifying the *mNeon* coding region using the primers neonF 5’-GCAAACAGTGAGCAAGGGCGAGGAGGATAAC-3’ and neonR 5’-GAGCTCGAGGTCGACTTACTTGT-3’ (Supplementary Table 6), and TOPO-TA cloning to pCR4 (Invitrogen). Depicted whole mount *in situ* results are representative of at least three biological replicates.

### Flow cytometry

Single cell suspensions were prepared from trunks of embryos anesthetized in 0.04% tricaine (MS-222). Trunks cells were manually dissociated by trituration in 2 mg/ml collagenase (Sigma) in 1X APBS (100 mM NaCl, 2.7 mM KCl, 2.8 mM KH_2_PO_4_, 8 mM Na_2_HPO_4_) at room temperature and monitored visually using a stereomicroscope. 4 ml of FACS buffer (1X APBS, 5% FBS (heat inactivated), 0.5% BSA (tissue culture grade), 10 U/ml heparin) was added and the cells were filtered through a 40 µm filter and washed twice in FACS buffer followed by centrifugation for 5 minutes at 1000 g. Cells were maintained on ice and promptly processed for cell sorting. Depicted flow cytometry results are representative of at least three biological replicates.

Fluorescent protein expression was measured on a five-laser LSR Fortessa flow cytometer (Becton Dickinson); GFP was excited by a 488 nm laser, and mCherry by a 561 nm laser. Cell doublets were sequentially excluded from the gating strategy based on both forward and side scatter properties. For cell sorting, populations of cells expressing GFP, mCherry, or both, were isolated using a Sony SH800 cells sorter (100 µm chip). Doublets were excluded based on scatter. FACS data were analysed using the FACSDiva software (BD Biosciences).

### Light and confocal microscopy

Embryos were photographed in 3% methyl cellulose on a Leica M205-FA stereomicroscope. Confocal images were acquired at the indicated magnifications with 0.2-0.8 µm z-steps on a Nikon C2 laser scanning confocal microscope as previously described^11^. Time lapse imaging was as previously described^11^ using the Nikon Elements 4.30 software. Experiments are representative of at least three biological replicates.

### RT-PCR

RT-PCR was performed as described previously^48^ using the primers and annealing temperatures indicated in Supplementary Table 6. RT-PCR results depicted in Supplementary Data Fig. 2 are representative of three independent experiments.

### Morpholinos

The Twist1bMO, Twist2MO, and Pax9MO were previously described^34,35^ and were injected at Twist1bMO 6 ng, Twist2MO 4 ng and Pax9MO 6 ng. A new splice-blocking Pax1aMO (5’-CTAAAACACTTTGAGTCTCACCTGT-3’) targeting the exon 1a / intron 1a boundary (exon 1 *pax1a-201* splice donor; exon 2 *pax1a-202* splice donor) was designed and validated for the working dose of 4 ng by RT-PCR (Supplementary Fig. 3).

## Data Availability

Data supporting the findings in this study, in particular details of the novel transgenic line *Tg(en.pf1-2.5pax1a:Neon)^sj6^* and the novel mutant lines *pax1a^sj7^* and *twist2^sj8^*, have been deposited with and will be publicly available at the Zebrafish Information Network (ZFIN; http://zfin.org/).

## Acknowledgments

The authors wish to acknowledge D. Traver and G. Crump for transgenic and mutant lines. P. Mead and the St. Jude flow cytometry core for FACS assistance, J. Peters, V. Frohlich, and the St. Jude imaging core for imaging assistance, and E. Damm for technical assistance. This work was supported by an NHLBI Loan Repayment Award and NIH F32HL129819 to N.O.G., European Research Council (ERC) Consolidator Grant-Rendox (ERC-CoG 647057) to M.M.S., and MOD BOC #5-FY-14-42, NIH R00HL097150, and NIH R01DK113973 to W.K.C.

## Author Contributions

The work was conceived and planned by W.C., C.K., and N.G. Experiments were performed by C.K. and N.G. The manuscript was written by W.C. and revised and edited with C.K., N.G., and M.S. The *acta2:mCherry* animals were generated by D.G. and M.S.

## Supplementary Information

Supplementary Information is available for this paper.

