## Supplementary Figures for "Sclerotome-derived vascular smooth muscle progenitors contribute to the haematopoietic stem cell specification niche"

#### Supplementary Figure Legends

**Supplementary Figure 1. Construction and validation of the *pax1a:Neon* transgenic line.** Schematic of the *pax1a* genomic locus (a) on Chromosome 17 from 17:42416407-42278340 (GRCz11), with the 1.9 kb putative promoter fragment, coding exons, and *pf1* enhancer regions indicated. Schematic representation of the alternative initiation sites used by transcripts *pax1a-201* (blue) and *pax1a-202* (yellow), with overlapping regions merged as green. Schematic of the transgenesis targeting construct with additional indication of the heterologous *SV40 polyA* and Tol2 transposase cis targeting elements indicated (c). Single color *in situ* comparison (d-g) of the expression of endogenous *pax1a* transcript (d, f) and transgenically expressed *Neon* transcript (e, g) and double *in situ* comparison (h-k) of *pax1a* (blue) and *Neon* (red-brown) at 23 hpf. Although the pharyngeal endoderm (dashed box region d-g, h, j) expresses endogenous *pax1a*, the transgene is not expressed there. Lateral views, dorsal up, anterior left (d, e, h, i). Dorsal views, anterior left (f, g, j, k). 40X (d-f, h, j) or 80X (i, k).

**Supplementary Figure 2. RT-PCR gene expression analysis of trunk endothelial cells.** *Kdrl:mCherry; fli1a:EGFP* double transgenic embryos were sorted by flow cytometry for single and double positive populations and analysed by RT-PCR for the indicated genes. Whole embryos and samples lacking reverse transcriptase (-RT) were analysed as controls.

**Supplementary Figure 3. Validation of the Pax1aMO.** Schematic of the exon structure of alternatively initiated *pax1a-201* (blue) and *pax1a-202* (yellow) transcripts, with overlapping regions in green. The region encoding the paired-box DNA-binding domain is depicted in olive drab. Location of the exon/intron boundary targeted by the morpholino (red bar) and primers (1F and 1Rb; Supplementary Table 6) used to analyse splice alterations in the morphants are indicated. Wild type, intron-trapped, and cryptic splice variants are schematically depicted at lower left with species sizes, and RT-PCR analysis displaying the sequence-validated wild type (Uninj lane) and splice-variant species (pax1aMO lane) with sizes indicated at lower right.

**Supplementary Figure 4. Neural crest migrate normally in Pax1aMO and Pax9MO animals.** Whole mount *in situ* depicting normal migration of neural crest cells labeled by *crestin* at 28 hpf (a, b) and 30 hpf (c, d) in uninjected (uninj) vs. Pax1aMO (b) or vs. Pax9MO (d). Lateral 80X views, dorsal up, anterior left.

**Supplementary Figure 5. Generation of a *pax1a* mutant.** Schematic of the exon structure of alternatively initiated *pax1a-201* (blue) and *pax1a-202* (yellow) transcripts, with overlapping regions in green. UTR in light shades, coding regions in dark shades. Location of the regions targeted by gRNA#7 and gRNA#2 are indicated (red arrows), as well as genotyping primer binding sites (F3b, F3c, R3; Supplementary Table 6). CRISPR/Cas9-mediated deletion of a 2.6 kb fragment results in the sequencing-validated locus depicted, and permits identification of the F3c/R3 wild-type (500 bp) or F3b/R3 mutant (300 bp) bands by genomic multiplex PCR depicted bottom right.

**Supplementary Figure 6. Generation of a *twist2* mutant.** Schematic of the exon structure of the *twist2* locus, containing a 5' exon containing 5'-UTR (open box), the complete coding region (blue), and partial 3'-UTR (open box), and a downstream 3'-UTR exon (open box). The region encoding the bHLH DNA-binding domain is indicated (grey). Location of the regions targeted by gRNA#3 and gRNA#9 are indicated (red arrows) as are the location of twist2 F and twist2 R primers used to identify mutant alleles (Supplementary Table 6). CRISPR/Cas9 injection yields 21 bp and 372 bp deletions within the coding region as indicated and verified by genomic PCR and sequencing.

**Supplementary Figure 7. Genetic compensation in the *pax1a*, *twist1b*, and *twist2* mutants.** Normal expression of *runx1* in wild type embryos (a, e, i) is comparable to that observed in the *pax1a* (b), *twist1b* (f), and *twist2* (j) mutants (verified by PCR genotyping). Although Pax1aMO, Twist1bMO, and Twist2MO are effective in wild type and heterozygous embryos (c, g, k), they are no longer effective in homozygous mutant embryos (d, h, l), indicating that the mutant displays compensatory gene expression. Blue arrowheads indicate regions of *runx1*$^{+}$ HSC precursors. Lateral views, dorsal up, anterior left, 80X.

**Supplementary Table Legends**

**Supplementary Table 1. Number of embryos displaying sclerotome alteration phenotypes.** The number of uninjected, Pax1aMO, Pax9MO, Twist1bMO, and Twist2MO injected embryos displaying reduced or otherwise altered expression as noted for the sclerotome markers *pax1a*, *pax9*, *twist1b*, and *twist2* are presented. n.d. indicates analysis not performed. Associated with Fig. 3.

**Supplementary Table 2. Number of embryos displaying control gene expression phenotypes.** The number of uninjected, Pax1aMO, Pax9MO, Twist1bMO, and Twist2MO injected embryos displaying altered gene expression as indicated for myotome (*myod1*), primitive erythrocyte (*gata1a*), endothelium (*cdh5*), arterial specification (*efnb2a*), and Notch receptor expression (*notch1b*) are presented. Associated with Fig. 5.

**Supplementary Table 3. Number of embryos displaying unaltered neural crest patterning.** The number of uninjected, Pax1aMO, and Pax9MO injected embryos displaying alterations in neural crest patterning are presented. Associated with Supplementary Fig. 4.

**Supplementary Table 4. Number of embryos displaying reduce markers of definitive haematopoiesis.** The number of uninjected, Pax1aMO, and Pax9MO injected embryos displaying normal or reduced levels of *runx1*, *cmyb*, and *rag1* as indicated. Associated with Fig. 6. Note that the results of discrete experiments with multiple biological replicates analysing effects on *runx1* are presented in Supplementary Tables 4 and 5.

**Supplementary Table 5. Genetic compensation in *pax1a*, *twist1b*, and *twist2* mutants.** The number of embryos expressing medium to high levels of *runx1* or reduced to absent levels of *runx1* are presented for wild type, *pax1a*, *twist1b*, and *twist2* mutant embryos comparing the effects of MO injection in wild type or correlated mutant are presented. Associated with Supplementary Fig. 7. Note that the results of discrete experiments with multiple biological replicates analysing effects on *runx1* are presented in Supplementary Tables 4 and 5.

**Supplementary Table 6. Primers used in this study.** Primers used for cloning, genotyping, and RT-PCR are listed with the associated primer Tms.

**Supplementary Movie Legends**

**Supplementary Movie 1. Three-pane timelapse of *pax1a:*Neon+ sclerotome migration from 20 - 30 hpf.** Schematics of the region depicted in the movie appear in Fig. 1e, j. Lateral, anterior left, dorsal up single-z view of confocal timelapse of *pax1a:*Neon+ cells in comparison to *kdrl:*mCherry+ endothelium and bright field from 20 - 30 hours (bottom) in the mid-trunk region of the developing embryo. The white bounding box indicates where the lateral, anterior left, dorsal up max-projection (top right) and virtual transverse reconstruction (top left, dorsal up) are extracted from. The DA, containing the haemogenic endothelium is visible as the upper red vessel, especially in the transverse section. Top movies 160X; lower movie 100X.

**Supplementary Movie 2.** A schematic of the region depicted appears in Fig. 1e. Lateral, anterior left, dorsal up, single-z view of confocal time lapse of *pax1a:*Neon+ cells in comparison to *kdrl:*mCherry+ endothelium and bright field from 20 - 30 hours in the mid-trunk region of the developing embryo. The white bounding box indicates the region depicted in Extended Movie 3. Same as lower pane of Supplementary Movie 1. 160X.

**Supplementary Movie 3.** A schematic of the region depicted appears in Fig. 1e. Lateral, anterior left, dorsal up max-projection view of confocal timelapse of *pax1a:*Neon+ cells in comparison to *kdrl:*mCherry+ endothelium and bright field from 20 - 30 hours in the mid-trunk region of the developing embryo. Same as upper right pane of Supplementary Movie 1. 160X.

**Supplementary Movie 4.** A schematic of the region depicted appears in Fig. 1j. Virtual transverse reconstruction (dorsal up) of confocal timelapse of *pax1a:*Neon+ cells in comparison to *kdrl:*mCherry+ endothelium and bright field from 20 - 30 hours in the mid-trunk region of the developing embryo. Neon+ (green) sclerotome cells migrate upwards and in between the PCV (lower red vessel) and DA (upper red vessel). Same movie as upper left pane of Supplementary Movie 1. 160X.

Supplementary Fig. 1

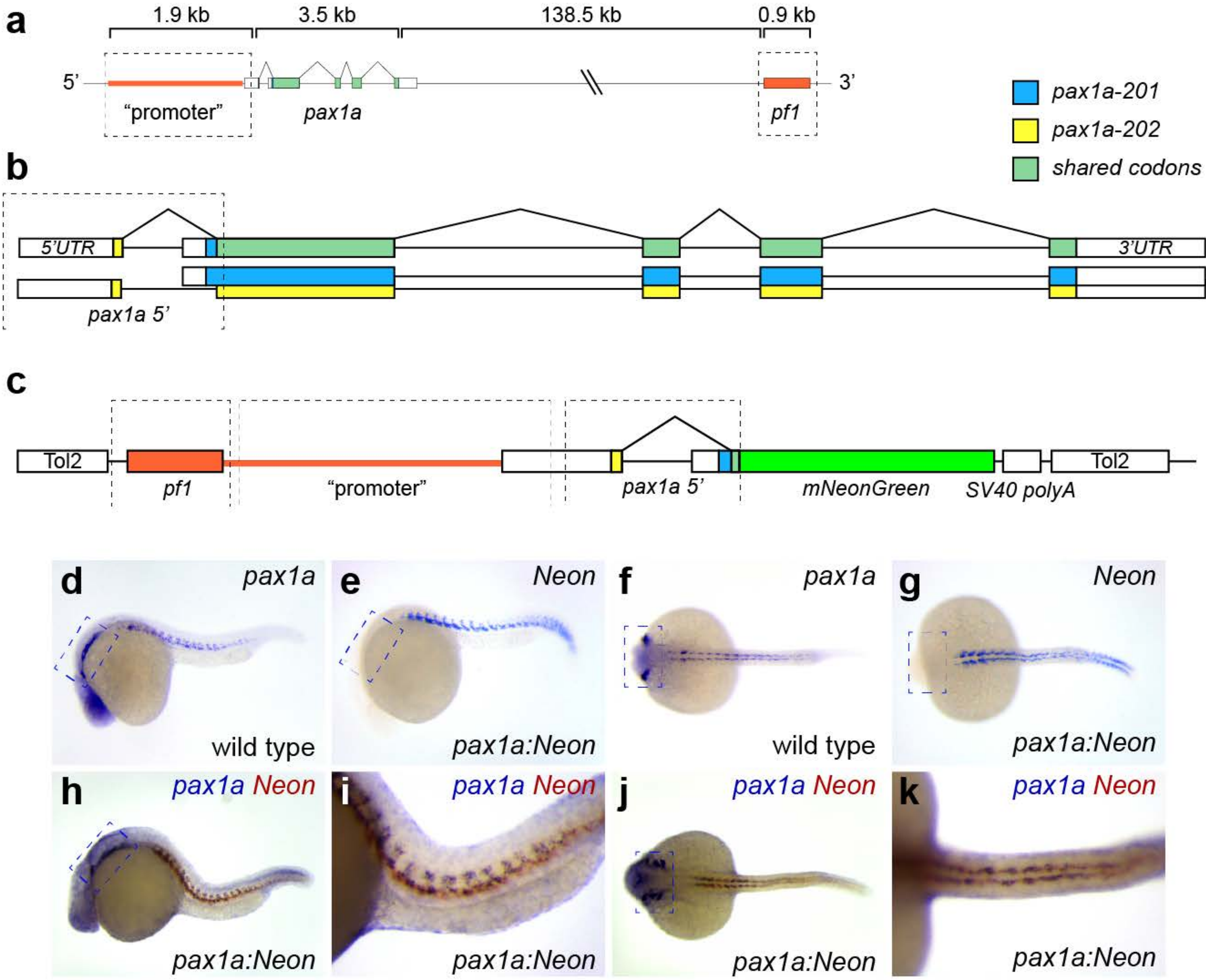

Supplementary Fig. 2

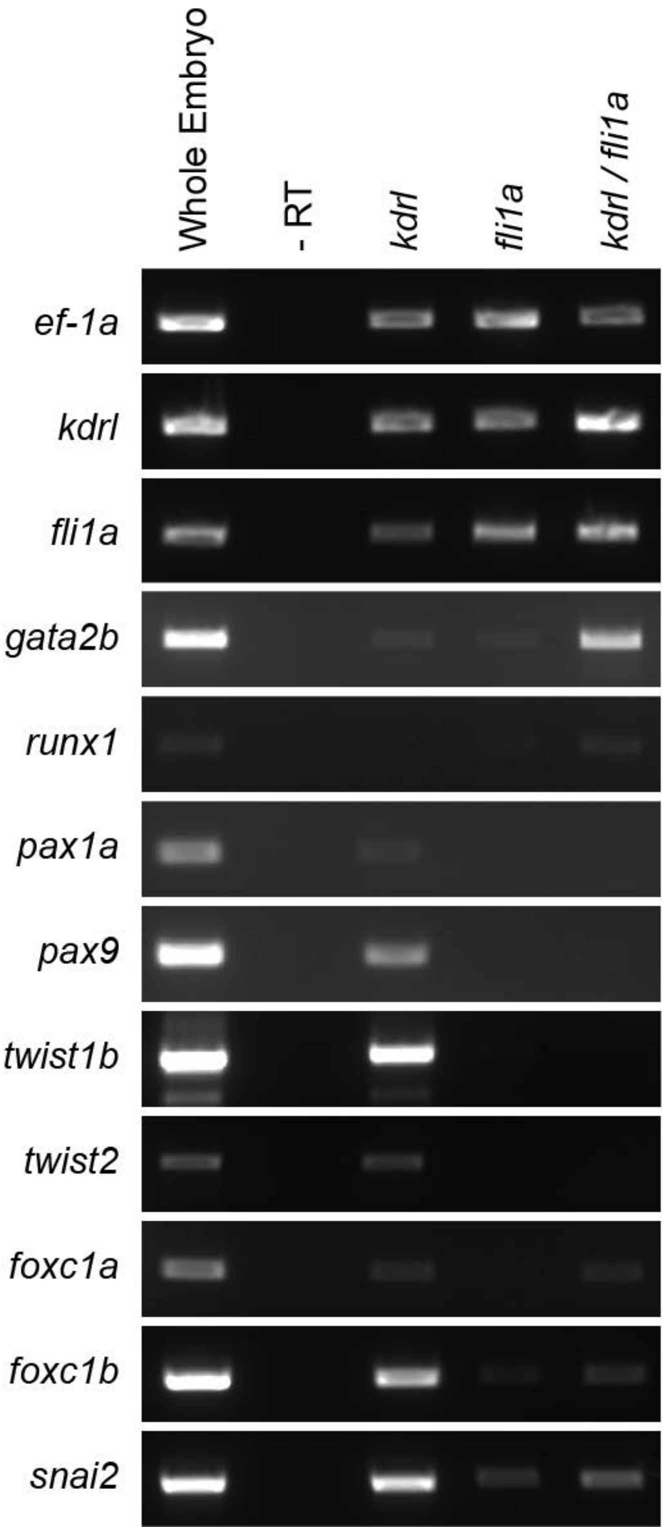

### Supplementary Fig. 3

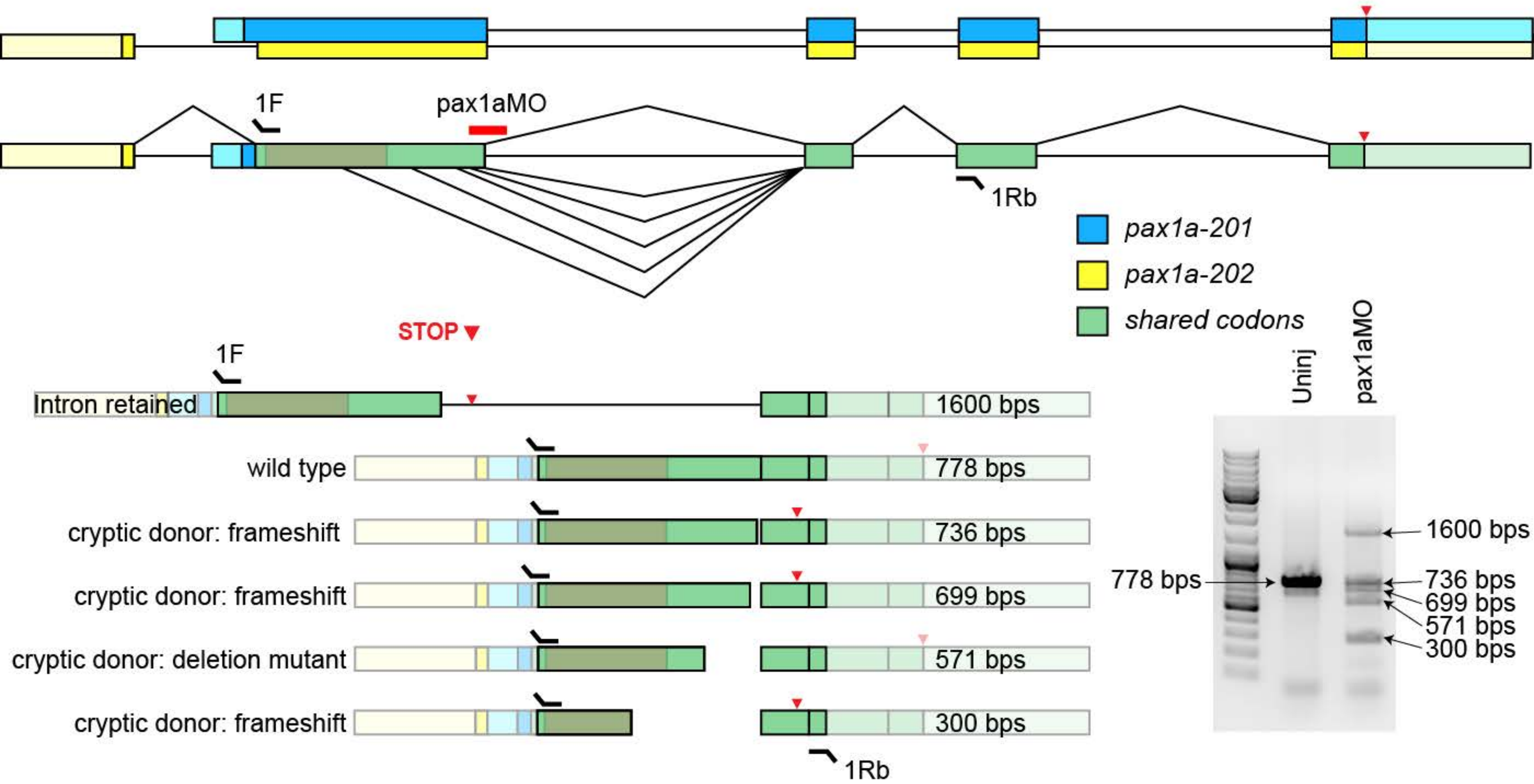

Supplementary Fig. 4

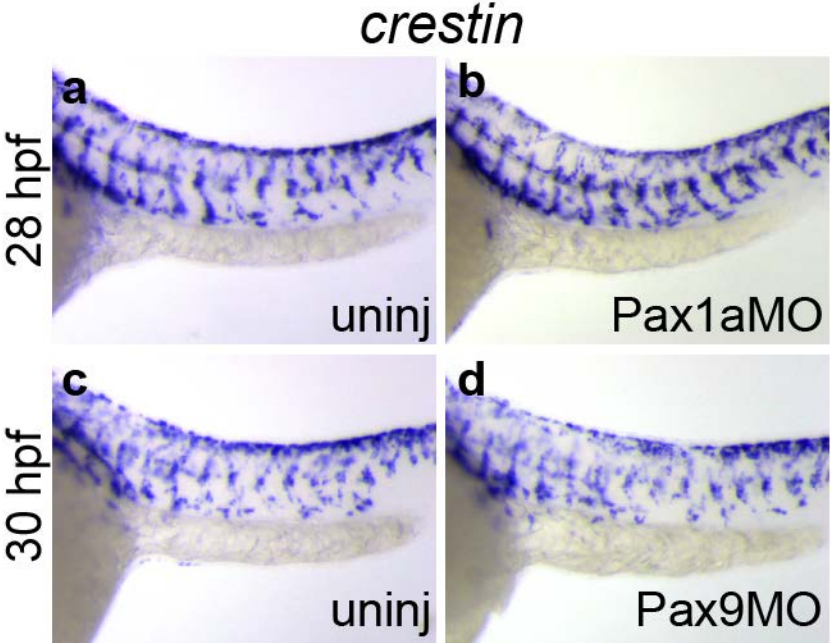

### Supplementary Fig. 5

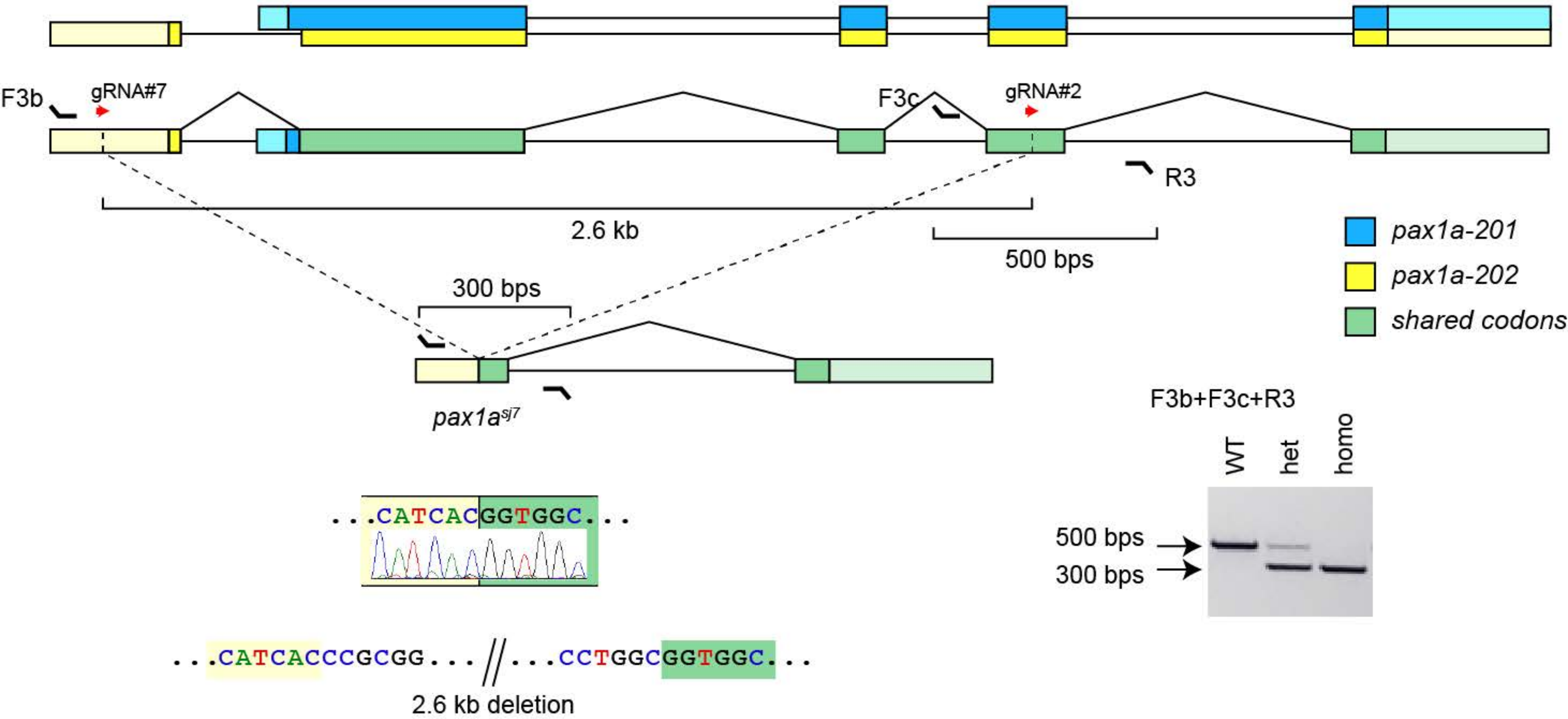

### Supplementary Fig. 6

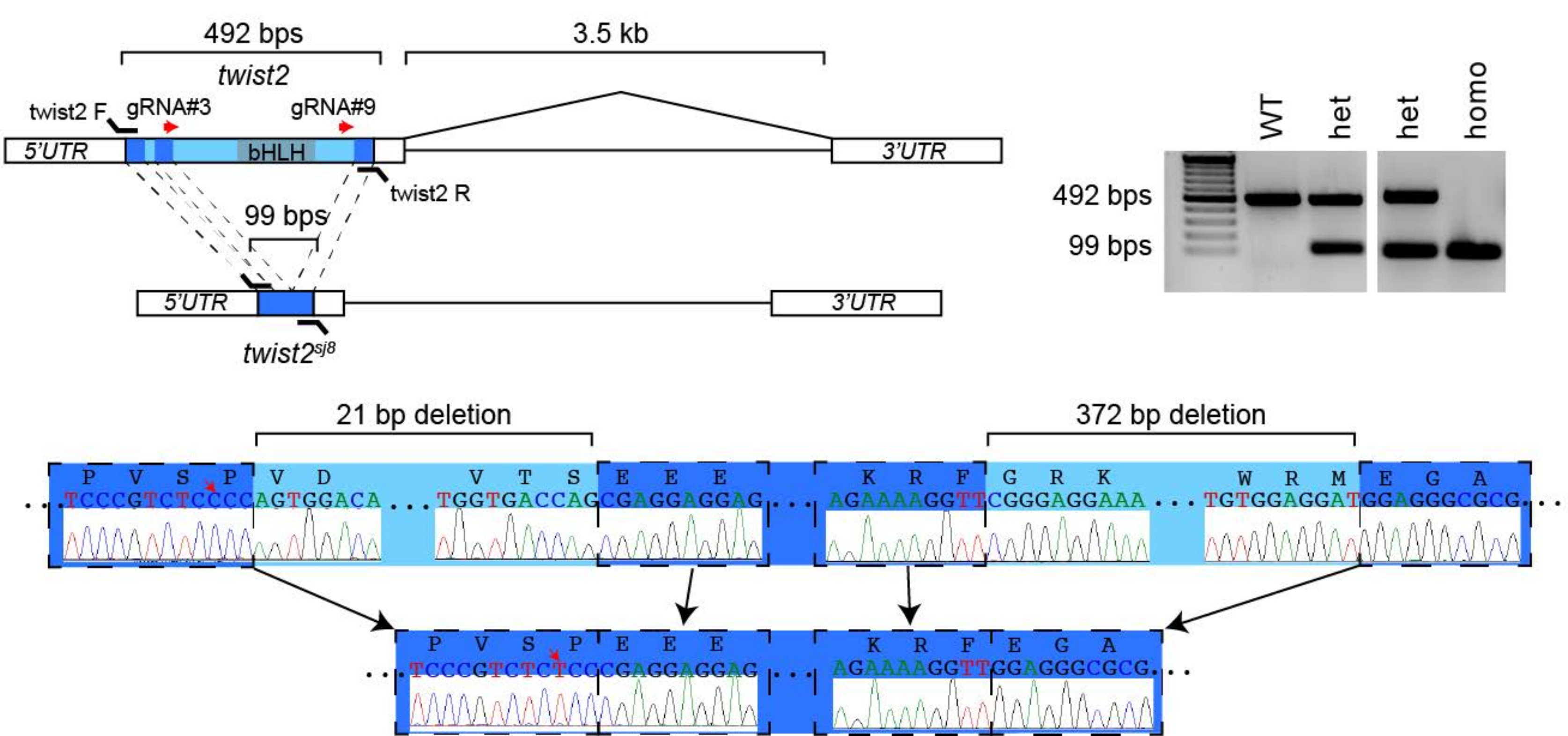

**Supplementary Data Table 1**

| *pax1a* | normal | reduced/absent |
|---|---|---|
| Uninj | 130 | 27 |
| Pax1aMO | n.d. | n.d |
| Pax9MO | 10 | 65 |
| Twist1bMO | 2 | 82 |
| Twist2MO | 16 | 58 |

| *pax9* | normal | reduced/absent |
|---|---|---|
| Uninj | 166 | 8 |
| Pax1aMO | 2 | 43 |
| Pax9MO | n.d. | n.d. |
| Twist1bMO | 0 | 76 |
| Twist2MO | 34 | 113 |

| *twist1b* | normal | reduced/absent |
|---|---|---|
| Uninj | 79 | 33 |
| Pax1aMO | 4 | 35 |
| Pax9MO | 11 | 58 |
| Twist1bMO | n.d. | n.d. |
| Twist2MO | 7 | 69 |

| *twist2* | normal | reduced/absent |
|---|---|---|
| Uninj | 149 | 16 |
| Pax1aMO | 3 | 60 |
| Pax9MO | 1 | 58 |
| Twist1bMO | 0 | 49* |
| Twist2MO | n.d. | n.d. |

*same pattern, reduced intensity

**Supplementary Data Table 2**

| *myod1* | normal | fuzzy/diffuse |
|---|---|---|
| Uninj | 108 | 4 |
| Pax1aMO | 44 | 6 |
| Pax9MO | 38 | 6 |
| Twist1bMO | 8 | 0 |
| Twist2MO | 17 | 0 |

| *gata1a* | normal | reduced/absent |
|---|---|---|
| Uninj | 124 | 17 |
| Pax1aMO | 44 | 20 |
| Pax9MO | 30 | 5 |
| Twist1bMO | 12 | 0 |
| Twist2MO | 11 | 0 |

| *cdh5* | normal | reduced/absent |
|---|---|---|
| Uninj | 146 | 4 |
| Pax1aMO | 54 | 0 |
| Pax9MO | 28 | 0 |
| Twist1bMO | 19 | 0 |
| Twist2MO | 17 | 4 |

| *efnb2a* | normal | reduced/absent |
|---|---|---|
| Uninj | 123 | 12 |
| Pax1aMO | 54 | 9 |
| Pax9MO | 50 | 7 |
| Twist1bMO | 24 | 0 |
| Twist2MO | 18 | 0 |

| *notch1b* | normal | reduced/absent |
|---|---|---|
| UI Tot | 151 | 9 |
| Pax1aMO | 43 | 0 |
| Pax9MO | 50 | 7 |
| Twist1bMO | 20 | 0 |
| Twist2MO | 25 | 0 |

**Supplementary Data Table 3**

| *crestin* | normal | slightly reduced |
|---|---|---|
| Uninj | 110 | 5 |
| Pax1aMO | 30 | 5 |
| Pax9MO | 51 | 3 |

**Supplementary Data Table 4**

| *runx1* | normal | reduced/absent |
|---|---|---|
| Uninj | 372 | 127 |
| Pax1aMO | 16 | 137 |
| Pax9MO | 21 | 82 |
| Twist1bMO | 15 | 79 |
| Twist2MO | 5 | 70 |

| *cmyb* | normal | reduced/absent |
|---|---|---|
| Uninj | 326 | 67 |
| Pax1aMO | 7 | 70 |
| Pax9MO | 19 | 47 |
| Twist1bMO | 5 | 44 |
| Twist2MO | 10 | 36 |

| *rag1* | normal | reduced/absent |
|---|---|---|
| Uninj | 153 | 94 |
| Pax1aMO | 1 | 61 |
| Pax9MO | 3 | 38 |
| Twist1bMO | 0 | 28 |
| Twist2MO | 10 | 36 |

**Supplementary Data Table 4**

| *runx1* | normal | reduced/absent |
|---|---|---|
| WT | 236 | 49 |
| *pax1a* mt | 71 | 16 |
| WT+Pax1aMO | 91 | 104 |
| mt+Pax1aMO | 43 | 16 |
| WT | 105 | 11 |
| *twist1b* mt | 61 | 18 |
| WT+Twist1b MO | 50 | 144 |
| mt+Twist1bMO | 55 | 33 |
| WT | 101 | 19 |
| *twist2* mt | 66 | 46 |
| WT+Twist2 MO | 11 | 78 |
| mt+Twist2MO | 36 | 19 |

#### Supplementary Data Table 5

| Purpose | Primer | Sequence (5'-3') | Tm |
|---|---|---|---|
| Cloning | neonF | GCAAACAGTGAGCAAGGGCGAGGAGGATAAC | 62 |
| | neonR | GAGCTCGAGGTCGACTTACTTGT | 62 |
| | pax9F | TCTAGAATGGAGCCAGCCTTT | 62 |
| | pax9R | ATGGATCCTCATAGAGCTGAAGCCACCAG | 62 |

| | | | |
|---|---|---|---|
| Genotyping | P1aCCF3b | CCCATTGCTCTCCGCATAAC | 60 |
| | P1aCCR3 | AGGCCACTGTTGAGATCATGC | 60 |
| | P1aCCF3c | GCCAAGCACGGATAATGTGT | 60 |
| | twist2 F | ATGGAAGAGAGTTCTAGCTCTCCC | 59 |
| | twist2 R | CTAGTGGGACGCAGACATCG | 59 |

| | | | |
|---|---|---|---|
| RT-PCR | pax1a For 1F | AGCAAACATACGGGGAGGTG | 60 |
| | pax1a rev | CGTAACTCGTGGTTTGCTGC | 60 |
| | pax1a 1Rb | GCTAGACAGGGTCGACGAAG | 60 |
| | pax9 for | GCAAACAGGCCTCTCATCCT | 60 |
| | pax9 rev | CTTCTAGTTTGGCGCTGGGA | 60 |
| | gata2b for | GAGTATCCCGCCAGTGTGTT | 60 |
| | gata2b rev | CTGTTGCGTGTCTGAATACC | 60 |
| | runx1 for | TCTGAGCAGTTGAGGCGAAG | 60 |
| | runx1 rev | GCAGGTATGTGTGGTAGCGT | 60 |
| | kdrl for | TGTATCCACCGTGATCTGGC | 60 |
| | kdrl rev | TCGGCAGCAGAATTCCTCAT | 60 |
| | fli1a for | CGGGCTCCACTGAAAATTGC | 60 |
| | fli1a rev | CCCGCCCACCATTTTATTGC | 60 |
| | twist1b for | AGCTCGACCCTTGGGAAAAG | 60 |
| | twist1b rev | ACAGCCATGACCTCTGTTGG | 60 |
| | twist2 for | CGCTCCCCTCGGATAAACTC | 60 |
| | twist2 rev | TGACAATACTCGCAGGTAGCC | 60 |
| | snai2 for | ACCCAAGACTCTGAGCTGGT | 60 |
| | snai2 rev | CACTTTGCACTTGTTCCCCG | 60 |
| | foxc1a for | CAATCCAGAACTCCCCGGAC | 60 |
| | foxc1a rev | TCCTTGTCCTTCATGGCGTC | 60 |
| | foxc1b for | AGCATGTACTCTCACGCCAC | 60 |
| | foxc1b rev | GCACCTTCACAAAGCACTCG | 60 |
| | ef1a for | CTTCTCAGGCTGACTGTGC | 60 |
| | ef1a rev | CCGCTAGCATTACCCTCC | 60 |
